# Individual Level Differential Expression Analysis for Single Cell RNA-seq data

**DOI:** 10.1101/2021.05.10.443350

**Authors:** Mengqi Zhang, Si Liu, Zhen Miao, Fang Han, Raphael Gottardo, Wei Sun

## Abstract

Bulk RNA-seq data quantify the expression of a gene in an individual by one number (e.g., fragment count). In contrast, single cell RNA-seq (scRNA-seq) data provide much richer information: the distribution of gene expression across many cells. To assess differential expression across individuals using scRNA-seq data, a straightforward solution is to create “pseudo” bulk RNA-seq data by adding up the fragment counts of a gene across cells for each individual, and then apply methods designed for differential expression using bulk RNA-seq data. This pseudo-bulk solution reduces the distribution of gene expression across cells to a single number and thus loses a good amount of information. We propose to assess differential expression using the gene expression distribution measured by cell level data. We find denoising cell level data can substantially improve the power of this approach. We apply our method, named IDEAS (Individual level Differential Expression Analysis for scRNA-seq), to study the gene expression difference between autism subjects and controls. We find neurogranin-expressing neurons harbor a high proportion of differentially expressed genes, and ERBB signals in microglia are associated with autism.

## Background

Single cell RNA-seq (scRNA-seq) data provide an unprecedented high-resolution view of gene expression variation within a bulk tissue sample, and thus help improve our understanding of the molecular basis of complex human diseases. For example, by comparing scRNA-seq data between cases and controls, we may identify cell-type-specific gene expression signatures that are related to disease etiology and progression [1, 2].

Early scRNA-seq studies often collect many cells from one or a few individuals and seek to compare gene expression between two groups of cells. Several methods have been developed towards this end [3–7]. As the scRNA-seq techniques evolve from a new revolution to a standard approach, many researchers start to collect scRNA-seq data from multiple individuals, and thus differential expression (DE) testing across individuals becomes an imperative task. The existing cell level DE methods are inappropriate for individual level DE testing as the sampling space of the cell level DE methods are cells but not individuals, and a significant p-value asserts DE if we sample more cells from the **same** set of individuals. In contrast, we focus on statistical inference in the population, i.e., whether we observe DE if we collect scRNA-seq data from more individuals.

In this paper, we assume the cells have been clustered into a few cell types if needed, and then we assess DE for each cell type separately. For individual level DE, one may first estimate cell type-specific gene expression per individual by pooling the gene expression across cells of the same cell type to create pseudo-bulk RNA-seq data for each cell type and then apply DE testing methods for bulk RNA-seq data [8, 9]. This pseudo-bulk approach captures shift of mean expression but may miss higher-order differential expression patterns, e.g., variance changes. To fully exploit the information in scRNA-seq data, we propose a new approach that captures the cell type-specific gene expression of an individual by a probability distribution and then compare such distributions across individuals. We refer to our method as Individual level <u>D</u>ifferential <u>E</u>xpression <u>A</u>nalysis for <u>S</u>cRNA-seq data (IDEAS).

## Results

### An overview of IDEAS

IDEAS performs DE testing gene by gene with respect to a categorical or continuous variable. To simplify the discussion, we consider a simple situation of two-group comparison between cases and controls (Figure 1). The first step of IDEAS is to estimate the distribution of one gene’s expression in each individual using a parametric or non-parametric method, conditioning on cell level covariates. The parametric method can be estimating a negative binomial (NB) or zero-inflated negative binomial (ZINB) distribution. The non-parametric method can be kernel density estimation or empirical estimation of cumulative distribution function (CDF). The next step is to calculate the distance between the gene expression distributions of any two individuals by the Jensen-Shannon divergence (JSD, using density estimates) or Wasserstein distance (Was, using CDF estimates) [10]. The final step is to assess whether within-group distances tend to be smaller than between-group distances. We define our test statistics by a pseudo F-statistic [11] and its null distribution can be estimated by permutations. This permutation procedure is computationally efficient because we do not need to re-calculate the distance matrix for each permutation. When sample size is large (e.g., n > 50), based on the connection between distance-based regression and kernel regression [12], we can use the asymptotic results from kernel regression to calculate p-values [13]. More details of IDEAS method are presented in the Methods Section.

**Figure 1.**
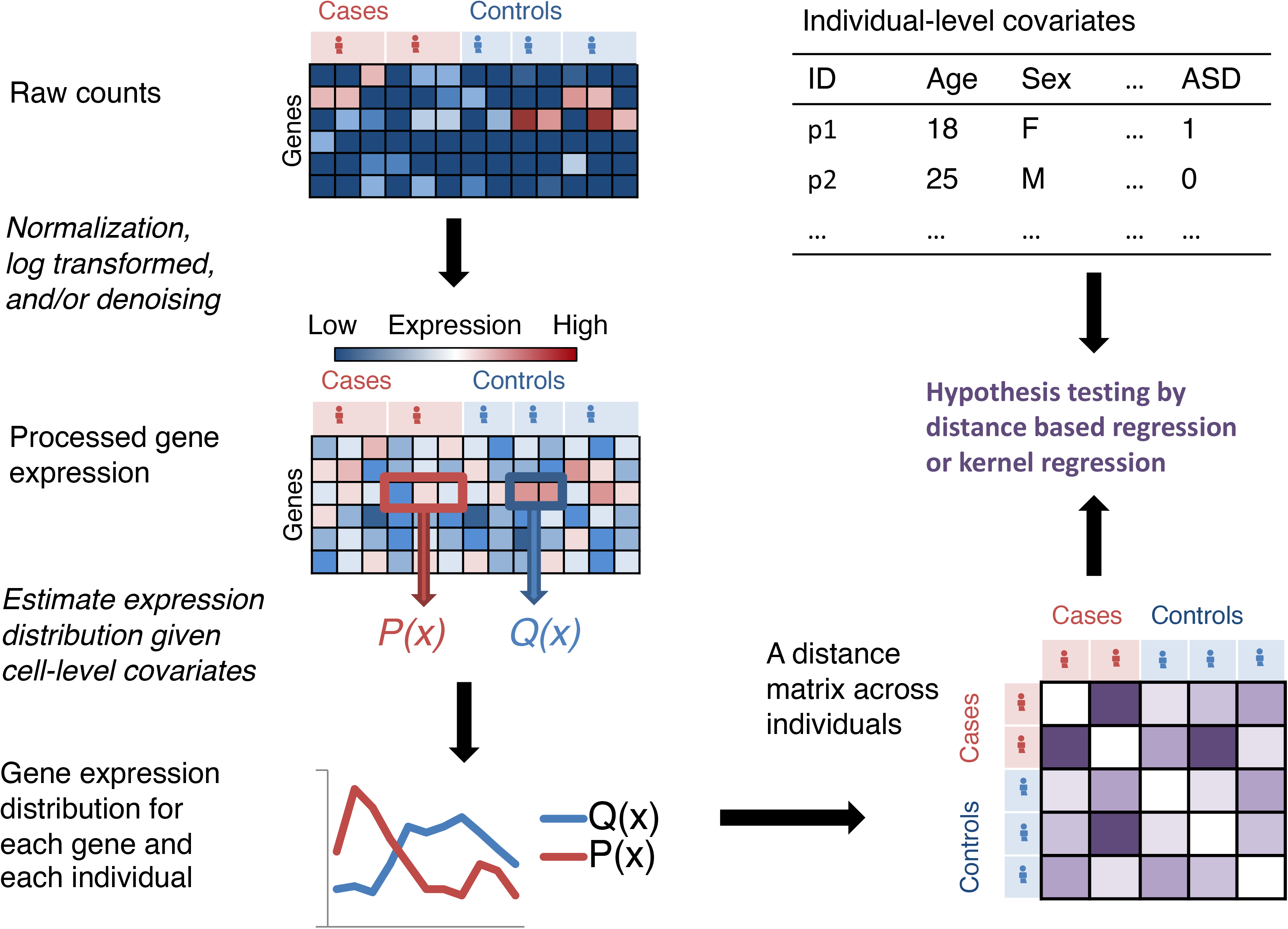
An overview of the IDEAS pipeline.

### Design of simulation studies

We evaluated the performance of IDEAS as well as a pseudo-bulk approach using simulations. We used DESeq2 [9] for DE analysis on pseudo-bulk data that were generated by adding up the counts across all the cells per individual. We considered four versions of our methods, with two methods to estimate within-individual distributions (ZINB or non-parametric (NP)) and two methods to estimate distances across individuals (JSD or Was).

We simulated scRNA-seq data based on a real dataset of 62,166 cells from the prefrontal cortex (PFC) of 13 autism patients and 10 controls [1] in the following steps. First, we estimated a ZINB distribution for each gene and each cell using a data denoising neural network method called DCA (deep count autoencoder) [14]. Next, we focused on the 8,626 L2/3 neuron cells to guide our simulation. We simulated the expression for 8,000 genes in 23 individuals (13 cases and 10 controls, 360 cells per individual) with one-to-one correspondence to the 8,000 genes that were expressed in the highest fractions of the cells (roughly > 20% of the 8,626 L2/3 neuron cells). For each gene, we assumed a ZINB distribution for its expression in the *i*-th individual and estimated four parameters: *µ_i_ =* log(mean), *φ_i_ =* log(over-dispersion), *π_i_ =* logit(proportion of zero-inflation), and *σ_i_* (the log-transformed standard deviation of log(mean) across all the cells of the *i*-th individual). The first three parameters were estimated by taking median over the cell level estimates by DCA. We estimated a multivariate normal distribution for these four parameters across the 23 individuals. Finally, we used this distribution to simulate parameters for 23 individuals and used the simulated parameters to simulate count data from ZINB distributions. We divided the 8,000 simulated genes into three groups. In the first and second 1,000 genes, we added DE signal in mean value (meanDE, 1.2-fold change) and variance (varDE, 1.5 fold change), respectively. The remaining 6,000 genes did not have any DE signal, and were referred to as equivalently expressed (EE) genes. These EE genes were used to evaluate the type I error.

### IDEAS can identify more DE patterns than the pseudo-bulk method

DESeq2 has slightly inflated type I error, slightly higher power than IDEAS for meanDE situation, and almost no power in the varDE situation (Figure 2A). In contrast, all four IDEAS methods control type I error very well and they all have much higher power than DESeq2 in the varDE situation (Figure 2A). We also illustrate a varDE gene that shows no DE signal in pseudo-bulk data (Figure 2B) while the difference of variation can be detected when examining the distribution of gene expression (Figure 2C).

**Figure 2.**
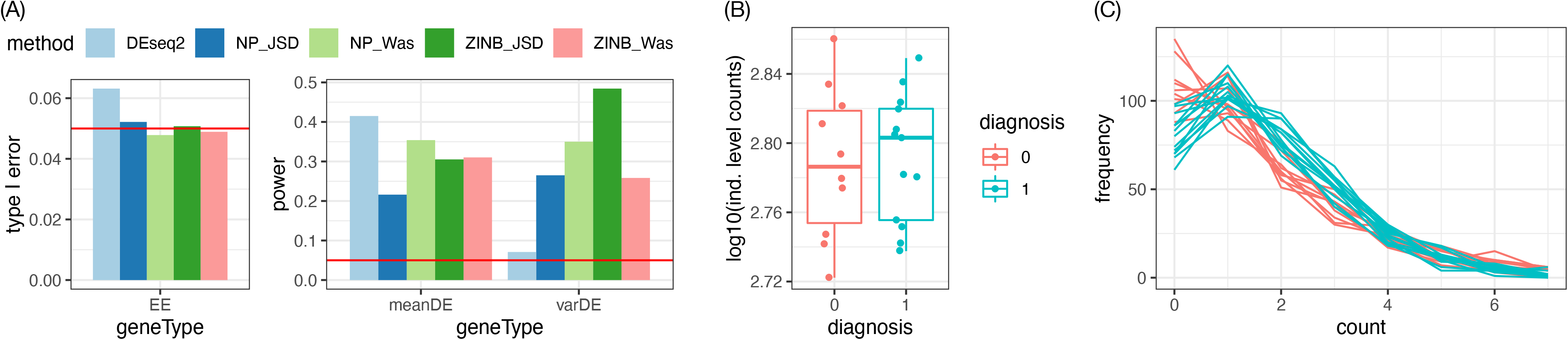
**(A)** Simulation results for 6,000 equivalently expressed (EE) genes, 1,000 genes with DE signal in mean value, and 1,000 genes with DE signal in variance. The distribution of gene expression within each individual was estimated by non-parametric (NP: kernel density for the Jensen-Shannon divergence (JSD) and empirical CDF for the Wasserstein distance (Was)) or parametric (ZINB: zero-inated negative binomial) method, respectively. **(B)** Illustration of the pseudo-bulk data of 23 individuals for one gene with DE signal on variance. **(C)** Illustration of the same gene in panel **(B)** for its empirical distribution in 23 individuals. The counts are truncated at 7 for illustration.

NB is sufficient to capture gene expression distribution derived from UMI counts. Several recent studies have shown that a NB distribution is often sufficient to model the scRNA-seq data using UMI (unique molecular identifier) [15–18]. When applying IDEAS on simulated data, the results (the p-values for all the genes) using NB or ZINB distribution are highly consistent (Supplementary Figure 1). Comparison of NB versus ZINB distribution using real data reaches similar conclusions (Supplementary Figure 2). Therefore, by default, NB distribution is used in the following analysis. Our implementation still allows ZINB distribution, which may be useful for scRNA-seq data generated without using UMI.

### Parametric approach is more robust to the sparsity of the scRNA-seq data

The cell level read-depth often varies considerably (Figure 3A), and thus needs to be accounted for when estimating individual-specific distributions. Adjusting for cell level read-depth (or any other cell level covariates) is straightforward for the parametric approach. We can run a NB regression against the log-transformed read-depth and then use the conditional NB distribution when setting the read-depth to certain value (e.g., the median value from all the cells across individuals). For the non-parametric approach, we can employ a linear regression with log-transformed counts as the response variable and the log-transformed read-depth as a covariate, and then use the fitted value given median read-depth plus residuals as the read-depth corrected data.

**Figure 3.**
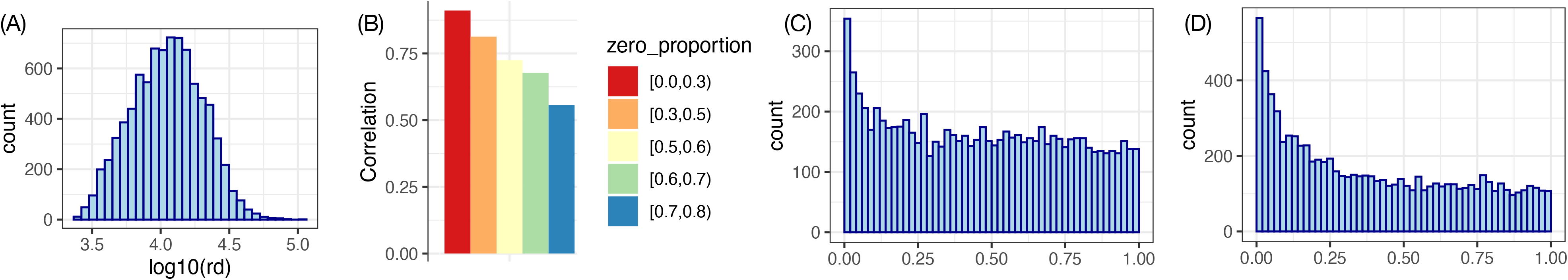
**(A)** Distribution of cell-level read-depth, with median around 10,000 and range from 2,414 to 105,488. **(B)** Correlation between the -log10(p-values) of DE testing between the two approaches to estimating individual-specific distribution: ZINB or empirical CDF. The genes were divided into 5 groups based on the proporion of cells where the observed gene expression is 0. **(C)** The P-value distribution when estimating individual-specific distribution using ZINB, followed by a distance calculation using the Wasserstein distance and a p-value calculation using permutation. **(D)** Same as (C) except that the input data are not the observed counts but the counts sampled from the cell specific ZINB estimated by DCA [14].

At this stage, the shortcoming of non-parametric method becomes obvious. In the real dataset, after selecting the 8,260 genes that are expressed in at least 20% of 8,626 L2/3 neuron cells, there are still more than half of the genes with zero expression in more than 50% of the cells. In addition, the remaining non-zero counts tend to be small, e.g., 1 to 5. A linear regression with such sparse data is highly unreliable. We illustrate this by comparing the -log10(DE p-value) for autism subjects versus controls obtained by two approaches: NB fit or empirical CDF fit followed by a distance calculation using the Wasserstein distance. The genes are divided into 5 categories based on the proportion of 0’s across those 8,626 L2/3 neuron cells. The correlation of the two approaches decreases as the proportion of zero’s increases (Figure 3B). After manual examination and comparison with results from pseudo bulk approaches, we conclude this is mainly due to the limitation of non-parametric approaches to handle sparse count data. Therefore, in the following analysis we focus on distribution estimation by the parametric approach (i.e., estimation of an NB distribution). When calculating the distances between individuals, using JSD or Wasserstein distance does not make much difference. We choose to focus on the Wasserstein distance because of its optimal theoretical properties [10, 19].

### Data denoising improves the statistical power

One angle to explain the difference between IDEAS and the pseudo-bulk method is through the bias-variance trade-off. The pseudo-bulk method summarizes the expression of many cells by summing them up, which leads to information loss (potential bias) but reduced variance. On the other hand, IDEAS tries to harvest the information from individual cells at the cost of a potentially higher uncertainty in estimating the gene expression distribution across cells. One direction to improve IDEAS is via denoising the scRNA-seq data, which is a well-studied topic. A popular denoising method named DCA [14] is used in this paper. DCA estimates a ZINB distribution for each gene and each cell based on a low-dimensional space that can filter out some noise in the data.

We applied DCA to 62,166 cells of 17 cell types from the prefrontal cortex (PFC). To assess the consequence of DCA denoising, we first focused on the 8,626 L2/3 neuron cells, one of the most abundant cell types. We sampled 5 counts from each cell-specific ZINB estimated by DCA and pooled them across cells to estimate an NB for each individual. We then proceeded testing using the Wasserstein distance. This new approach using DCA denoised and augmented data (Figure 3D) has higher power than the same NB-Wasserstein approach using observed count data (Figure 3C).

This approach to sampling counts from cell-specific ZINB estimates by DCA is flexible since we can use the sampled counts to fit an NB regression to account for any cell level covariates. However, it is also computationally intensive. An alternative approach is to directly estimate individual-level distributions by averaging the cell specific ZINBs estimated by DCA (Supplementary Materials Section 1.1). This direct computation approach gives similar results to the results from NB regression (Supplementary Materials Section 2.2). Therefore, we use this direct computation approach in the following analysis.

### IDEAS combined with denoising improves the power to identify DE genes

We performed DE analysis between autism subjects and controls for all 17 cell types. We only considered the genes that were expressed in at least 20% of the cells for each cell type, and the number of genes varied a lot across cell types: ranging from 578 (Microglia) to 9,291 (L5_6-CC) (Figure 4A). When controlling false discovery rate to be 10%, DESeq2, IDEAS, and IDEAS combined with DCA identified 268, 41, and 4,571 DE genes respectively (Supplementary Table 1). Most of the DE genes found by any of the three methods were from a few cell types: the interneuron cells and excitatory neurons on layer 2/3, as well as a subtype of neurogranin (NRGN)-expressing neurons (Neu-NRGN-II), suggesting the relevance of these cell types in autism.

**Figure 4.**
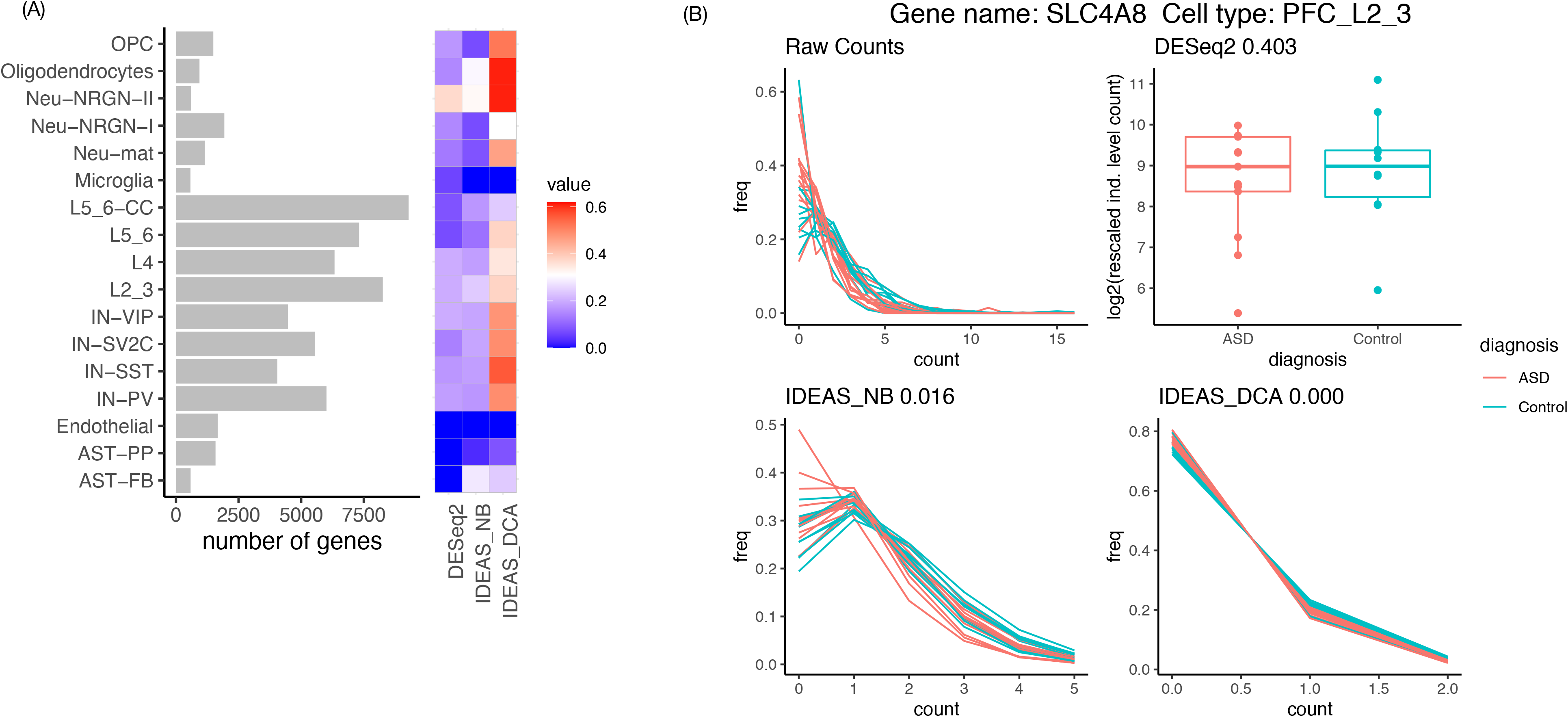
**(A)** The left panel shows the number of genes we studied for each cell type. A gene is included in our study if it is expressed in at least 20% of the cells. The right panel shows the estimates of the proportion of genes that are differentially expressed between autism subjects and controls for each cell type. **(B)** An example where IDEAS (combined with DCA) identifies strong DE signals while DESeq2 does not.

These results confirm that denoising scRNA-seq data can substantially improve the power of IDEAS. An example where the DE patterns become cleaner after denoising is shown in Figure 4B. This gene, SLC4A8, transports sodium and ions across cell membrane and is associated with glutamate release by neurons [20], and thus it can have functional role in autism development.

We further estimated the proportion of DE genes using a p-value distribution [21]. Here we estimated the proportion of DE genes as 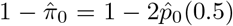, where 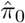 was the estimated proportion of non-DE genes and 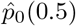 was the proportion of genes with p-values larger than 0.5. The DE proportion estimates were higher for excitatory neurons (e.g., layer 2/3 or layer 4 excitatory neurons) and interneurons (e.g., vasoactive intestinal polypeptide (VIP) and somatostatin (SST) expressing interneurons), partly due to relatively higher expression level in these cell types. In contrast, few DE signals were detected in astrocytes, endothelial cells, or microglia, possibly due to low gene expression in these cell types. Neu-NRGN-II was again an exception where the number of expressed genes was low while all three methods identified high proportion of DE genes (Figure 4A, Supplementary Table 2). Neurogranin (NRGN) is a calmodulin-binding protein, and it has been associated with Alzheimer’s disease [22] and schizophrenia [23]. Our results suggest that it is also potentially associated with autism.

### IDEAS improves the power to identify autism-related genes

The Simons Foundation Autism Research Initiative (SFARI) has compiled a list of autism risk genes. Most of these genes are identified because they harbor more disruptive mutations in autism subjects than in a general population. DNA mutations cannot directly affect biological function. At least part of their effect on biological systems is mediated through gene expression, and thus these genes may be identified by DE analysis. We assessed whether there is significant overlap between cell type-specific DE genes (which are defined using a liberal p-value cutoff of 0.05) and SFARI genes. IDEAS combined with DCA identified significant overlaps in four cell types: excitatory neurons on layer 2/3 or layer 4, and interneurons expressing SST or VIP (Figure 5A). In contrast, neither IDEAS nor DESeq2 identifies any significant overlap. We observe similar patterns if we just ask whether SAFRI genes tend to have smaller p-values by gene set enrichment analysis (GSEA) (Supplementary Table 3). Lack of significant overlap between SFARI genes and DE genes could indicate small proportions of overlap or limited power to identify DE genes and thus that overlap is not statistically significant. For example, small proportion of overlap is the main reason for excitatory neurons on layer 2/3 (L2 3) (Figures 5B and 5D) while limited power to identify DE genes is the main reason for interneurons expressing VIP (IN-VIP) (Figures 5C and 5E).

**Figure 5.**
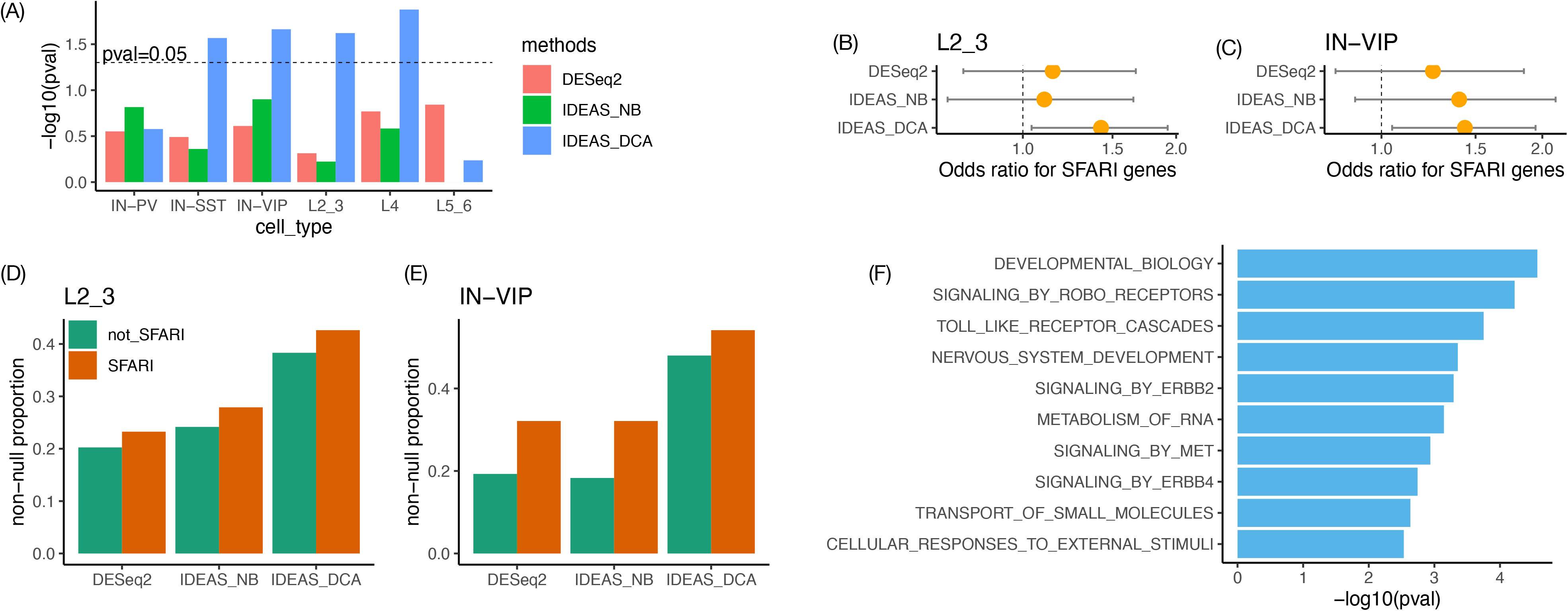
**(A)** Fisher’s exact test p-values to assess whether autism related genes (SFARI genes) have significant overlap with differential expressed genes (nominal p-values < 0.05). **(B-C)** Odds ratios and their 95% confidence intervals derived from Fisher’s exact test in panel (A) for two cell types: layer 2/3 excitatory neurons (L2 3) and vasoactive intestinal polypeptide (VIP){expressing interneurons (IN-VIP). **(D-E)** Estimates of the proportion of DE genes among those SFARI genes or non-SFARI genes for two cell types: L2 3 and IN-VIP. **(F)** Pathways that are over-represented by the genes that are differentially expressed between autism subjects and controls in microglia.

### Differentially expressed pathways in microglia

Microglia is of particular importance for autism because it is the resident immune cells in brain and immune response is an important factor of autism etiology [24]. In our analysis, although all the methods find few DE genes in microglia, gene set enrichment analysis (GSEA) that uses the ranking of all the genes identifies a few pathways that are differentially expressed between autism subjects and controls. At adjusted p-value cutoff of 0.05, GSEA using the DE ranking by DESeq2 identifies one pathway: signaling by ERBB4. Using the ranking by IDEAS combined with DCA, GSEA identifies this pathway together with 9 others, including signaling by ERBB2 (Supplementary Materials Section 2.5). An earlier study has shown that ERBB signals can lead to proliferation and activation of microglia [25]. Separate studies have also shown that exonic deletion of ERBB4 is associated with intellectual disability or epilepsy [26]. Our findings, combined with these earlier studies, suggest that ERBB signals may be an underlying factor that leads to different microglia activities between autism subjects and controls.

## Discussion

Our method IDEAS is designed for individual level DE analysis using single cell RNA-seq data. IDEAS compares gene expression distribution across individuals, and thus it can identify any pattern of DE including shift of mean or variance. Such flexibility is important for scRNA-seq data because of the heterogeneity of cell populations. For example, we can divide all the cells from a brain sample to excitatory neurons, interneurons, and a few glia cell types such as astrocyte, microglia, oligodendrocyte, etc.· However, excitatory neurons and interneurons can be further divided into many smaller categories. Therefore, the DE signal may exist in a subset of the cells and IDEAS is more suitable to capture such subtle DE patterns than the pseudo-bulk method that mainly assesses shifts in mean expression.

Methods designed to assess DE across cells can be modified using a mixed effect model framework to account for cell-cell dependence within an individual, and to perform DE across individuals. For example, MAST has such an option [4]. However, such approaches capture both cell-level and individual level DE signals, and when sample size is small, the cell-level variance may dominate the DE signal. Therefore, it should be used with caution especially when sample size is small.

A key step of IDEAS is to estimate gene expression distribution for each individual. We have demonstrated that non-parametric estimates of gene expression distribution are often unreliable, especially for the genes with low expression. Therefore we recommend parametric approaches, e.g., estimating gene expression distribution for one individual by a negative binomial distribution, which is a Poisson-Gamma mixture. In addition to the parametric or non-parametric estimates, an alternative is a semi-parametric one using a Poisson mixture with a non-parametric mixing distribution. We pursued this approach in a separate work [19].

We have observed that denoising scRNAseq data can improve the power of IDEAS. However, this also creates some uncertainty in real data analysis as the effect of denoising depends on the methods as well as the cell populations. In general, a denoising method works well if there is a latent structure in the data that are not confounded with case/control status. For example, considering scRNA-seq data of many cells that can be grouped into a few cell types, then the latent structure in the data are cell type-specific gene expression. Therefore, we recommend running the denoising procedure for all the cells of different types. We have used DCA for denoising. DCA handles read-depth difference across cells by a simple approach: dividing the observed read counts by read-depth. There is room to improve it by making more flexible correction of the read depth, for example, through a conditional variational autoencoder [27]. We will explore such more flexible denoising methods in a future work.

## Methods

### IDEAS

#### Input and output

The input data for IDEAS include gene expression data (a matrix of scRNA-seq fragment counts per gene and per cell), the variable of interest (e.g., case-control status), together with two sets of covariates. One set is cell level covariates, such as read-depth per cell. The cell level covariates are used to estimate the gene expression distribution of each individual across all the cells. The other set is individual-level covariates such as age, gender, batch effect, etc.· The output of IDEAS is a permutation p-value for each gene under the null hypothesis that the expression of this gene is not associated the variable of interest, given the rest covariates.

#### Calculation of distance matrix across individuals

We examine two metrics to evaluate the distance between the gene expression distributions of two individuals. One is the Jensen-Shannon divergence (JSD) and the other one is the Wasserstein distance. For two probability distributions denoted by *P* and *Q*:

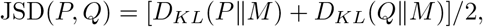

where *M* is a distribution whose density function is *f_M_* (*x*) = 0.5[*f_P_* (*x*) + *f_Q_*(*x*)], and 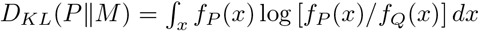 is the Kullback-Leibler divergence. The Wasserstein distance has attracted lots of attention in the machine learning fields recently [10]. We use the Wasserstein-1 distance, which is also referred to as the earth-moving distance. Intuitively, it is the minimum amount of effort to move the mass from one distribution to the other distribution. For one dimensional problem, the Wasserstein distance has a close form:

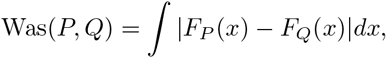

where *F_P_* (*x*) and *F_Q_*(*x*) denote the cumulative distribution functions for *P* and *Q*, respectively.

We explore two approaches to estimate the distribution of gene expression across all the cells of an individual. The first approach is a parametric one where we estimate the gene expression distribution by an NB or a ZINB distribution. Here we describe our method for the ZINB and NB is a special case for ZINB. Since our method is applied for each gene separately, we describe the procedure for one gene and ignore gene index to simplify the notation. Let *Y_i_* be a random variable for gene expression of individual *i*. Then a ZINB is a mixture distribution of a zero-inflation component and a negative binomial distribution component:

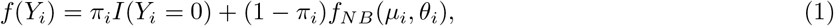

where *π_i_* is the zero-inflation proportion, *µ_i_* and *θ_i_* are the mean value and over-dispersion parameter for a negative binomial distribution, respectively, such that the variance of the negative binomial distribution is 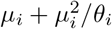.

Suppose we observe gene expression across *n_i_* cells of the *i*-th individual, denoted by **y***_i_*. A naive approach is to estimate a ZINB distribution using **y***_i_*. This approach will most likely have little power for differential expression testing because cell level read depth can vary a lot across cells / individuals, and thus it may dominate the estimated distribution and obscure any other signals. Therefore, we perform ZINB regression (or NB regression if NB distribution is assumed) of confounding factors such as the cell level read-depth.

The second approach is a non-parametric one. The specific solution depends on the distance metric used. For JSD, we estimate density using kernel method, by R function density with default Gaussian kernel. For Was, we use the R function wasserstein1d from R package transport, which takes input data points to calculate inverse of CDF and Wasserstein distance. In either case, we log transform the observed count data and then use a linear regression to obtain the adjusted log counts when all the covariates are set to their medians.

#### Data augmentation using auto-encoder

ScRNA-seq data are often noisy due to the limited number of RNA molecules per cell. Many methods have been developed to denoise scRNA-seq data. We employ one of such methods named deep count autoencoder network (DCA) [14] in our analysis. DCA exploits the low-dimensional structure of scRNA-seq data by an autoencoder, a neural network method. The input to DCA is the observed count matrix and the output is the estimated ZINB distribution for each gene and each cell. Note that this ZINB is a cell-specific distribution, and it is different from what we seek to estimate, which is an individual-specific distribution. We can achieve this goal in two ways. One is to simulate *m* counts from each cell-specific distribution and then pool them across cells to estimate the individual-specific distribution. This approach gives more flexibility to account for cell-level covariates though it is computationally intensive. The other approach is to directly add up the density estimates across cells to estimate the individual level distribution.

#### Calculation of p-value

We compare two approaches to calculate a p-value for each gene given the distance matrix across all individuals and individual level covariates. One is a distance-based test known as Permutational Multivariate Analysis of Variance (PERMANOVA) [11, 28], and the other one is kernel based regression implemented in R package MiRKAT [13]. When sample size is larger, kernel regression should have more computational advantages since it can calculate p-values using the asymptotic distribution of the test-statistic, though for studies with small or moderate sample sizes, kernel regression will also rely on permutation to assess p-values. For either kernel regression or PERMANOVA, the distance matrix *D* needs to be transformed to a kernel matrix **G** by

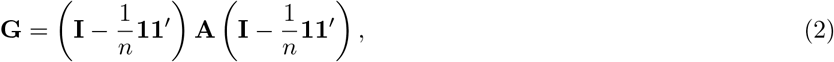

where **A =** *−*(1*/*2)**D**^2^ and **G** is the Grower’s centered matrix of **A** [11, 29]. This matrix may have some negative eigenvalues. Following earlier works, we set those negative eigenvalues to 0 [12, 30].

Let *Z* be the set of variables including the variable of interest (*X*) and all the covariates. Let **H***_Z_* be the hat matrix **H***_Z_ =* **Z**(**Z***^’^***Z**)^−1^**Z***^’^*, and denote the trace of a square matrix **U** by *tr*(**U**). Then the (pseudo) F statistic [11, 28] that quantifies the collective association between the gene expression and all the variables in *Z* is

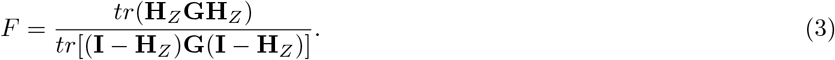

To generate the distribution of *F* under permutation, we permute *X* or re-sample *X* given all the covariates, following the PERMONVA-S method [31], and combine the permuted/re-sampled *X* with the covariates and generate a new data matrix *Z_p_*. The re-sampling approach is more desirable because it maintains the association between *X* and other covariates. However, when sample size is small, it could be unstable and thus we use permutation by default. Given *Z_p_*, we can calculate the F-statistic following equation (3). Note that our method is different from PERMONVA-S since we consider all the covariates when calculating the F-statistics while PERMONVA-S only considers the variable of interest. When the covariates have strong association with gene expression, our method can remove their impact and thus increases the power for testing.

## Availability of data and materials

The list of SFARI ASD risk genes was downloaded from https://gene.sfari.org/database/human-gene/. These genes were scored as “syndromic” (mutations that are associated with a substantial degree of increased risk and consistently linked to additional characteristics not required for an ASD diagnosis) and/or 7 categories from 1 to 7, with high, strong, and suggestive evidence for categories 1-3. Here we use the 350 genes that belong to the syndromic category or categories 1 to 7. Our software package and data analysis pipeline for simulations and autism data are available at https://github.com/Sun-lab/ideas.

## Competing interests

The authors declare that they have no competing interests.

**Consent for publication**

**Authors’ contributions**

