## supplementary materials for "Individual Level Differential Expression Analysis for Single Cell RNA-seq data"

April 29, 2021

### 1 Supplementary Methods

#### 1.1 Implementation of DCA-direct estimation method

DCA-direct method estimates the expression distribution of one gene across the cells of an individual by making use of the parameter estimates from DCA output in a straight forward manner. DCA takes the observed count matrix (rows for genes and columns for cells) as input and outputs four parameter matrices, normalized mean matrix $\bar{M}$, dispersion parameter matrix $\Theta$, dropout probability matrix $\Pi$, and mean matrix $M$. Each matrix has the same dimension as that of the input count matrix, with the value at each position providing a denoised parameter estimate for the corresponding gene and cell. The difference between normalized mean matrix $\bar{M}$ and mean matrix $M$ lies in DCA's adjustment for cell-level read-depth. To account for the factors that read-depth varies among different cells, DCA first normalizes the input raw count matrix by size factors computed based on cell-level read-depth, before proceeding to learn essential latent features and generate denoised parameter estimates. As an intermediate output of DCA for estimating the mean parameters, $\bar{M}$ is on the normalized scale. While as a final output, $M$ is on the original scale of raw count matrix, and it is derived by multiplying the size factors back to $\bar{M}$. For DCA-direct method, we make use of normalized mean matrix $\bar{M}$ together with $\Theta$ and $\Pi$.

For the simplicity of notation, we assume that the cells belong to one specific individual occupy the first $J$ columns of the count matrix. For one specific gene $i$ and a cell $j$, let $\pi_{ij}$, $\bar{\mu}_{ij}$ and $\theta_{ij}$ denote the corresponding elements on position $(i, j)$ of matrices $\Pi$, $\bar{M}$ and $\Theta$ respectively. The denoised distributions from DCA with normalized mean for the expression of this gene on the $J$ cells are:

$$P_{i1} = \text{ZINB}(\pi_{i1}, \bar{\mu}_{i1}, \theta_{i1}),\ P_{i2} = \text{ZINB}(\pi_{i2}, \bar{\mu}_{i2}, \theta_{i2}),\ \ldots,\ P_{iJ} = \text{ZINB}(\pi_{iJ}, \bar{\mu}_{iJ}, \theta_{iJ}).$$

For this individual, to get an expression distribution estimate for gene $i$ across cells, on each possible count value, we calculate the probability estimate simply by averaging the corresponding probabilities from $P_{ij}, j = 1, 2, \ldots, J$. Once we have estimated distributions for individuals, we can calculate the distance matrix either by Jensen-Shannon divergence or Wasserstein distance, and compute p-values through kernel-based association test or Permutational Multivariate Analysis of Variance.

### 2 Supplementary Results

#### 2.1 Comparison of NB vs. ZINB to characterize individual-level gene expression

In this section we compare p-values from using NB or ZINB distributions to characterize individual-level gene expression through running analysis on both simulated data and real count data. Figure 1 demonstrates the comparison on simulated data generated based on count data from the cell type excitatory neurons on layer 2/3 (L2/3) according to the process described in "Design of simulatin studies" subsection in main text. The p-values from these two approaches are very similar.

Figure 2 compares the p-values from IDEAS with NB and ZINB distributions on real count data from the cell type L2/3. The shape of the histograms and the scatter plot show that the results from these two approaches are highly consistent.

#### 2.2 Comparison of sampling-based approach vs. DCA-direct method

Besides DCA-direct method, another way to utilize DCA outputs is to generate count data according to the denoised distribution estimate given by DCA for each gene and each cell, then pool the data across cells and fit a negative distribution for each individual using the pooled data, calculate the distance matrix and compute p values. Following the notations as in the subsection "Implementation of DCA-direct estimation method" in main text, different from DCA-direct method, this DCA sampling-based approach makes use of the mean matrix $M$ instead of the normalized mean matrix $\bar{M}$, together with the dispersion parameter matrix $\Theta$ and dropout probability matrix $\Pi$.

Again for the simplicity of notation, we assume that the cells belonging to a specific individual occupy the first $J$ columns of the count matrix. For gene $i$ and cell $j$, let $\pi_{ij}$, $\mu_{ij}$ and $\theta_{ij}$ denote the corresponding elements on position $(i, j)$ of matrices $\Pi$, $M$ and $\Theta$ respectively. The denoised distributions from DCA with mean on the scale of original raw count for this gene on the $J$ cells are:

$$Q_{i1} = \text{ZINB}(\pi_{i1}, \mu_{i1}, \theta_{i1}),\ Q_{i2} = \text{ZINB}(\pi_{i2}, \mu_{i2}, \theta_{i2}),\ \ldots,\ Q_{iJ} = \text{ZINB}(\pi_{iJ}, \mu_{iJ}, \theta_{iJ}).$$

Then for each gene and each cell, we can generate multiple counts (for example, $m = 5$, 10, or 20) from the corresponding denoised distribution:

$$\begin{aligned} c_{i11}, c_{i12}, \ldots, c_{i1m} &\sim Q_{i1}, \\ c_{i21}, c_{i22}, \ldots, c_{i2m} &\sim Q_{i2}, \\ \ldots & \quad \ldots \\ c_{iJ1}, c_{iJ2}, \ldots, c_{iJm} &\sim Q_{iJ}. \end{aligned}$$

Now for this individual and gene $i$, we have in total $mJ$ counts sampled from the denoised distributions for the original $J$ cells. Next, we treat all these $mJ$ sampled counts as counts from different cells, and apply IDEAS on the matrix consisting of these sampled counts.

Figure 3 compares the p-values from DCA sampling-based and DCA-direct approaches, under generated sample size $m = 5, 10, 20$ for DCA sampling-based approach. The trends look relatively consistent, but the DCA sampling-based approach is computationally much more expensive due to the extra computational cost to fit NB distributions.

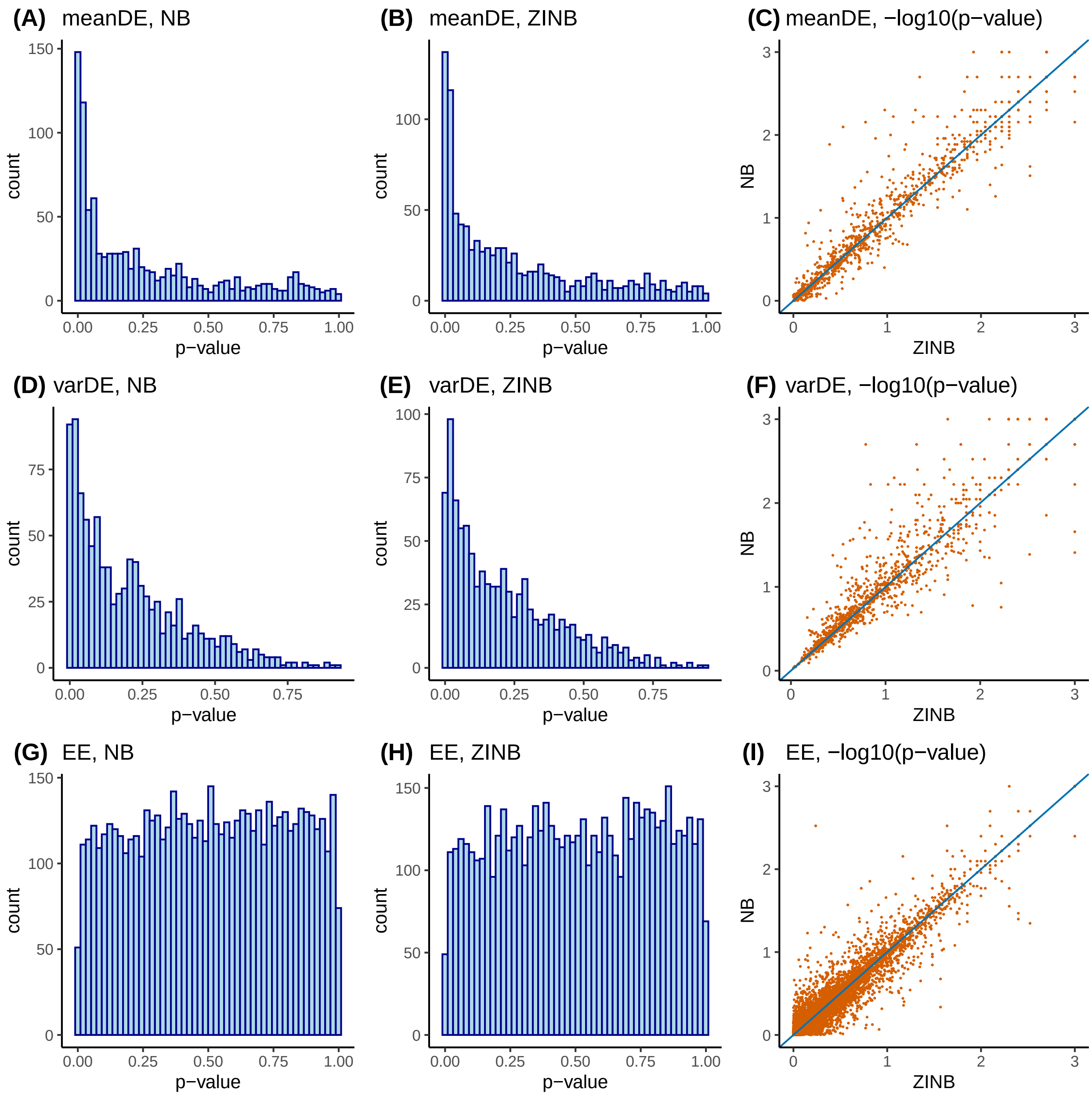

Figure 1: Comparison between the pvalues from NB and those from ZINB approaches on simulated data. Data generation process is described in "Design of simulation studies" subsection under the Result section of the main text. Each row corresponds to one group of simulated genes, with top, middle and bottom row corresponding to `meanDE`, `varDE` and `EE` respectively. The first column gives the histogram of p-values from using NB distribution in each group, and the second column gives those from using ZINB distribution. The last column provides the scatter plot comparing NB and ZINB approaches on their negative log10-transformed p-values. Each point in the scatter plot corresponds to one of the 8,000 simulated genes. For both NB and ZINB, the cell-level covariate to adjust for is read-depth, the method used to calculated distance between gene expression distribution is Wasserstein distance (Was). The p-values are computed through PERMANOVA.

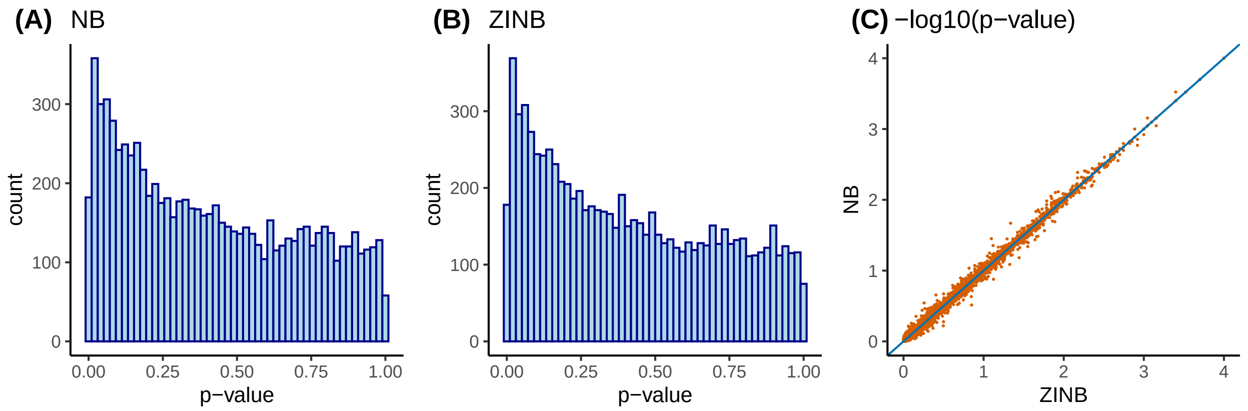

Figure 2: **(A-B)** Histogram of p-values from NB or ZINB approach on real data of cell type L2/3. **(C)** Scatter plot of -log10(p-values) from NB v.s. that from ZINB approach. Each point in the plot corresponds to one of the 8,260 genes expressed in at least 20% of 8,626 L2/3 neuron cells. The value on x-axis gives the negative log-transformed p-value with based 10 from ZINB, and the value on y-axis gives that from NB. For both NB and ZINB, the cell-level covariate to adjust for is read-depth, the method used to calculated distance between gene expression distribution is Wasserstein distance (Was), and the p-values are computed through PERMANOVA.

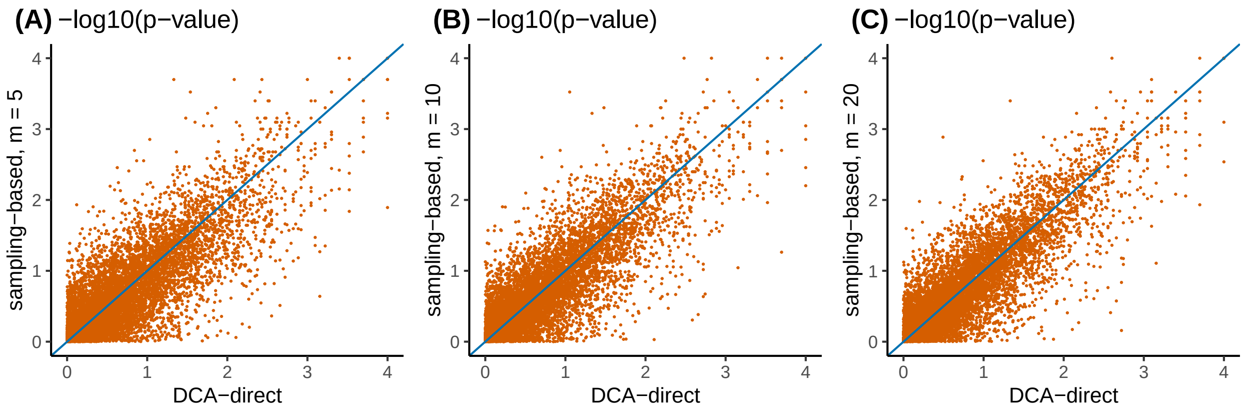

Figure 3: Scatter plots of -log10(p-values) from DCA sampling-based and DCA-direct approaches. Both approaches rely on the cell-level expression distributions estimated by DCA. The major difference is DCA sampling-based approach first sample cell-level counts and then estimate individual-level distributions, while DCA-direct approach directly estimates individual-level distributions by averaging the cell-level distributions. The cell type here is L2/3. **(A)** Comparison between -log10(p-values) from DCA sampling-based approach ($y$-axis) with $m = 5$ and those from DCA-direct approach ($x$-axis). **(B-C)** Similar comparison to that in (A), except that (B) has $m = 10$ and (C) has $m = 20$ for DCA sampling-based approach on $y$-axis. For DCA sampling-based approach, we fit a NB distribution per individual, with adjustment for the cell-level read-depth. For both DCA sampling-based and DCA-direct approaches, the method taken to calculated distance between gene expression distribution is Wasserstein distance (Was), and the p-values are computed through PERMANOVA.

#### 2.3 The number of DE genes

Table 1: The number of DE genes identified by different methods using FDR 0.1 as cutoff.

| Cell Type | number of genes | DESeq2 | IDEAS_NB | IDEAS_DCA |
|---|---|---|---|---|
| AST-FB | 584 | 0 | 0 | 0 |
| AST-PP | 1579 | 1 | 0 | 0 |
| Endothelial | 1665 | 1 | 0 | 0 |
| IN-PV | 6010 | 18 | 0 | 985 |
| IN-SST | 4049 | 60 | 5 | 1064 |
| IN-SV2C | 5555 | 6 | 0 | 1066 |
| IN-VIP | 4470 | 39 | 0 | 800 |
| L2_3 | 8260 | 15 | 0 | 260 |
| L4 | 6332 | 9 | 0 | 0 |
| L5_6 | 7313 | 0 | 0 | 0 |
| L5_6-CC | 9291 | 7 | 0 | 0 |
| Microglia | 578 | 0 | 0 | 0 |
| Neu-mat | 1154 | 2 | 0 | 173 |
| Neu-NRGN-I | 1930 | 25 | 9 | 0 |
| Neu-NRGN-II | 593 | 67 | 27 | 158 |
| Oligodendrocytes | 939 | 16 | 0 | 0 |
| OPC | 1490 | 2 | 0 | 65 |
| Total | 61792 | 268 | 41 | 4571 |

Table 2: Estimates of the proportion of DE genes

| **Cell Type** | **n_genes** | **DESeq2** | **IDEAS_NB** | **IDEAS_DCA** |
|---|---|---|---|---|
| AST-FB | 584 | 0 | 0.28 | 0.24 |
| AST-PP | 1579 | 0.004 | 0.03 | 0.083 |
| Endothelial | 1665 | 0 | 0 | 0.004 |
| IN-PV | 6010 | 0.177 | 0.172 | 0.488 |
| IN-SST | 4049 | 0.17 | 0.18 | 0.564 |
| IN-SV2C | 5555 | 0.141 | 0.189 | 0.492 |
| IN-VIP | 4470 | 0.199 | 0.189 | 0.483 |
| L2_3 | 8260 | 0.203 | 0.243 | 0.385 |
| L4 | 6332 | 0.196 | 0.18 | 0.348 |
| L5_6 | 7313 | 0.066 | 0.117 | 0.376 |
| L5_6-CC | 9291 | 0.075 | 0.166 | 0.238 |
| Microglia | 578 | 0.055 | 0 | 0 |
| Neu-mat | 1154 | 0.132 | 0.078 | 0.456 |
| Neu-NRGN-I | 1930 | 0.152 | 0.07 | 0.306 |
| Neu-NRGN-II | 593 | 0.373 | 0.323 | 0.605 |
| Oligodendrocytes | 939 | 0.154 | 0.299 | 0.606 |
| OPC | 1490 | 0.166 | 0.071 | 0.522 |

#### 2.4 Overlap with SAFRI genes

Table 3: Gene set enrichment analysis p-value for the enrichment of SAFRI genes based on rankings of DE signals

| | DESeq2 | IDEAS_NB | IDEAS_DCA |
|---|---|---|---|
| PFC_IN-PV | 0.09 | 0.27 | 0.18 |
| PFC_IN-SST | 0.11 | 0.54 | 0.032 |
| PFC_IN-VIP | 0.23 | 0.11 | 0.053 |
| PFC_L2_3 | 0.25 | 0.18 | 0.0045 |
| PFC_L4 | 0.074 | 0.091 | 0.032 |
| PFC_L5_6 | 0.049 | 0.061 | 0.98 |

#### 2.5 Gene set enrichment analysis (GSEA) using REACTOME pathways

We downloaded REACTOME pathway annotation `c2.cp.reactome.v7.1.symbols.gmt` from `https://data.broadinstitute.org/gsea-msigdb/msigdb/release/7.1/`, and ran GSEA using R package `fgsea` [1]. Here we list all the results with adjusted p-value smaller than 0.05.

```
---------------------------------------------------
Endothelial
---------------------------------------------------

$DESeq2
   pathway
 1: SRP_DEPENDENT_COTRANSLATIONAL_PROTEIN_TARGETING_TO_MEMBRANE
 2: TRANSLATION
 3: RRNA_PROCESSING
 4: RESPONSE_OF_EIF2AK4_GCN2_TO_AMINO_ACID_DEFICIENCY
 5: EUKARYOTIC_TRANSLATION_INITIATION
 6: SELENOAMINO_ACID_METABOLISM
 7: EUKARYOTIC_TRANSLATION_ELONGATION
 8: REGULATION_OF_EXPRESSION_OF_SLITS_AND_ROBOS
 9: NONSENSE_MEDIATED_DECAY_NMD
10: METABOLISM_OF_AMINO_ACIDS_AND_DERIVATIVES
11: INFLUENZA_INFECTION
12: INFECTIOUS_DISEASE
13: RESPIRATORY_ELECTRON_TRANSPORT_ATP_SYNTHESIS_AND_HEAT_PRODUCTION
14: METABOLISM_OF_RNA
            pval         padj      NES
 1: 3.352074e-06 0.0005296276 1.585903
 2: 3.213129e-06 0.0005296276 1.561339
 3: 1.185720e-05 0.0009367189 1.556261
 4: 1.045400e-05 0.0009367189 1.541682
 5: 1.565277e-05 0.0009892549 1.549457
 6: 2.326745e-05 0.0012254193 1.537243
 7: 8.373781e-05 0.0037801640 1.495820
 8: 1.008269e-04 0.0039826633 1.488167
 9: 1.820414e-04 0.0063916768 1.460114
10: 2.646323e-04 0.0083623813 1.441035
11: 7.168753e-04 0.0191212959 1.411776
12: 7.261252e-04 0.0191212959 1.310402
13: 1.812653e-03 0.0409141736 1.505900
```

```
14: 1.775471e-03 0.0409141736 1.278604

$IDEAS_DCA
   pathway
1: TRANSCRIPTIONAL_REGULATION_BY_TP53
           pval        padj      NES
1:  5.480576e-06 0.001731862 1.710584

---------------------------------------------------
IN-SST
---------------------------------------------------

$DESeq2
   pathway
1: RESPIRATORY_ELECTRON_TRANSPORT
2: RESPIRATORY_ELECTRON_TRANSPORT_ATP_SYNTHESIS_AND_HEAT_PRODUCTION
3: THE_CITRIC_ACID_TCA_CYCLE_AND_RESPIRATORY_ELECTRON_TRANSPORT
4: COMPLEX_I_BIOGENESIS
5: PEPTIDE_LIGAND_BINDING_RECEPTORS
           pval        padj      NES
1: 9.246451e-07 0.000370927 1.671747
2: 1.270298e-06 0.000370927 1.616130
3: 1.706289e-05 0.003321576 1.488195
4: 2.693380e-05 0.003932335 1.695798
5: 1.469840e-04 0.017167735 1.692564

$IDEAS_NB
   pathway                   pval       padj      NES
1: L1CAM_INTERACTIONS 6.323096e-05 0.03686365 1.486866

$IDEAS_DCA
   pathway
1: RESPIRATORY_ELECTRON_TRANSPORT
2: RESPIRATORY_ELECTRON_TRANSPORT_ATP_SYNTHESIS_AND_HEAT_PRODUCTION
3: THE_CITRIC_ACID_TCA_CYCLE_AND_RESPIRATORY_ELECTRON_TRANSPORT
4: RNA_POLYMERASE_II_TRANSCRIPTION
5: COMPLEX_I_BIOGENESIS
6: NOTCH_HLH_TRANSCRIPTION_PATHWAY
           pval        padj      NES
1: 8.208884e-06 0.002554043 1.731140
2: 9.050801e-06 0.002554043 1.681495
3: 1.312008e-05 0.002554043 1.580630
4: 1.454047e-04 0.021229087 1.288993
5: 2.600301e-04 0.030371520 1.724897
6: 4.353902e-04 0.042377977 1.783170

---------------------------------------------------
L2_3
---------------------------------------------------

$IDEAS_NB
   pathway                              pval       padj       NES
1: TRANSLESION_SYNTHESIS_BY_POLK 4.693707e-05 0.03783128 -2.264327
```

```
--------------------------------------------------
L5_6
--------------------------------------------------

$IDEAS_DCA
   pathway
1: RESPIRATORY_ELECTRON_TRANSPORT
2: THE_CITRIC_ACID_TCA_CYCLE_AND_RESPIRATORY_ELECTRON_TRANSPORT
3: RESPIRATORY_ELECTRON_TRANSPORT_ATP_SYNTHESIS_AND_HEAT_PRODUCTION
4: COMPLEX_I_BIOGENESIS
           pval        padj      NES
1: 3.874828e-06 0.002968119 1.559201
2: 3.032646e-05 0.011615034 1.427667
3: 4.903206e-05 0.012519519 1.453862
4: 1.293762e-04 0.024775540 1.553722

--------------------------------------------------
L5_6-CC
--------------------------------------------------

$DESeq2
   pathway                    pval     padj     NES
1: COMPLEX_I_BIOGENESIS 2.15753e-05 0.018339 1.57771

--------------------------------------------------
Microglia
--------------------------------------------------

$DESeq2
   pathway                    pval      padj      NES
1: SIGNALING_BY_ERBB4 0.0003435272 0.0340092 1.732156

$IDEAS_DCA
   pathway                                        pval        padj      NES
 1: DEVELOPMENTAL_BIOLOGY                 2.693380e-05 0.002666446 1.563263
 2: SIGNALING_BY_ROBO_RECEPTORS           5.983003e-05 0.002961586 1.823819
 3: TOLL_LIKE_RECEPTOR_CASCADES           1.774619e-04 0.005856243 1.792809
 4: SIGNALING_BY_ERBB2                    5.092329e-04 0.010082811 1.745213
 5: NERVOUS_SYSTEM_DEVELOPMENT            4.395162e-04 0.010082811 1.559152
 6: METABOLISM_OF_RNA                     7.145970e-04 0.011790851 1.693711
 7: SIGNALING_BY_MET                      1.151892e-03 0.016291038 1.705486
 8: SIGNALING_BY_ERBB4                    1.801664e-03 0.022295587 1.681338
 9: TRANSPORT_OF_SMALL_MOLECULES          2.320680e-03 0.025527478 1.554697
10: CELLULAR_RESPONSES_TO_EXTERNAL_STIMULI  2.924422e-03 0.028951778 1.590222

--------------------------------------------------
Neu-NRGN-I
--------------------------------------------------

$DESeq2
   pathway
1: RESPIRATORY_ELECTRON_TRANSPORT
2: RESPIRATORY_ELECTRON_TRANSPORT_ATP_SYNTHESIS_AND_HEAT_PRODUCTION
3: THE_CITRIC_ACID_TCA_CYCLE_AND_RESPIRATORY_ELECTRON_TRANSPORT
```

```
4: COMPLEX_I_BIOGENESIS
5: SIGNALING_BY_MODERATE_KINASE_ACTIVITY_BRAF_MUTANTS
           pval        padj      NES
1: 2.185732e-05 0.004163820 1.568409
2: 1.819099e-05 0.004163820 1.521486
3: 5.299993e-05 0.006730991 1.466350
4: 2.922455e-04 0.027836380 1.600551
5: 6.211871e-04 0.047334456 1.665379

$IDEAS_NB
   pathway
1: RESPIRATORY_ELECTRON_TRANSPORT
2: RESPIRATORY_ELECTRON_TRANSPORT_ATP_SYNTHESIS_AND_HEAT_PRODUCTION
3: COMPLEX_I_BIOGENESIS
4: THE_CITRIC_ACID_TCA_CYCLE_AND_RESPIRATORY_ELECTRON_TRANSPORT
5: SIGNALING_BY_MODERATE_KINASE_ACTIVITY_BRAF_MUTANTS
           pval        padj      NES
1: 3.444094e-07 0.000131220 1.668845
2: 6.597622e-06 0.001256847 1.560452
3: 1.045400e-05 0.001327658 1.701401
4: 2.383151e-05 0.002269951 1.489015
5: 2.651538e-04 0.020204723 1.687284

$IDEAS_DCA
   pathway
1: RESPIRATORY_ELECTRON_TRANSPORT
2: THE_CITRIC_ACID_TCA_CYCLE_AND_RESPIRATORY_ELECTRON_TRANSPORT
3: RESPIRATORY_ELECTRON_TRANSPORT_ATP_SYNTHESIS_AND_HEAT_PRODUCTION
4: COMPLEX_I_BIOGENESIS
           pval         padj      NES
1: 1.277296e-09 2.433250e-07 1.839651
2: 6.435037e-10 2.433250e-07 1.778831
3: 4.705457e-09 5.975930e-07 1.757933
4: 1.181424e-07 1.125307e-05 1.935357

--------------------------------------------------
Oligodendrocytes
--------------------------------------------------

$IDEAS_NB
   pathway                            pval       padj      NES
1: SIGNALING_BY_RHO_GTPASES 0.0003428697 0.04731602 1.632791
```

### References

[1] Gennady Korotkevich, Vladimir Sukhov, Nikolay Budin, Boris Shpak, Maxim N Artyomov, and Alexey Sergushichev. Fast gene set enrichment analysis. *BioRxiv*, page 060012, 2021.
